# Exploring sex-specific neuroendocrine influences on the sensorimotor-association axis in single individuals

**DOI:** 10.1101/2024.05.04.592501

**Authors:** Bianca Serio, Deniz Yilmaz, Laura Pritschet, Hannah Grotzinger, Emily G. Jacobs, Simon B. Eickhoff, Sofie L. Valk

**Affiliations:** Max Planck School of Cognition, Leipzig, Germany; Max Planck Institute for Human Cognitive and Brain Sciences, Leipzig, Germany; Institute of Neuroscience and Medicine, Brain & Behavior (INM-7), Research Centre Jülich, Jülich, Germany; Institute of Systems Neuroscience, Medical Faculty, Heinrich Heine University Düsseldorf, Düsseldorf, Germany; Palo Alto High School, Palo Alto, USA; Department of Psychological and Brain Sciences, University of California, Santa Barbara, USA

## Abstract

Human neuroimaging studies consistently show multimodal patterns of variability along a key principle of macroscale cortical organization – the sensorimotor-association (S-A) axis. However, little is known about day-to-day fluctuations in functional activity along this axis within an individual, including sex-specific neuroendocrine factors contributing to such transient changes. We leveraged data from two densely sampled healthy young adults, one female and one male, to investigate intra-individual daily variability along the S-A axis, which we computed as our measure of functional cortical organization by reducing the dimensionality of functional connectivity matrices. Daily variability was greatest in temporal limbic and ventral prefrontal regions in both participants, and was more strongly pronounced in the male subject. Next, we probed local and system-level effects of steroid hormones and self-reported perceived stress on functional organization. Our findings revealed modest effects that differed between participants, hinting at subtle –potentially sex-specific– associations between neuroendocrine fluctuations and intra-individual variability along the S-A axis. In sum, our study points to neuroendocrine factors as possible modulators of intra-individual variability in functional brain organization, highlighting the need for further research in larger samples.

## INTRODUCTION

Patterns of functional connectivity are considered to be broadly stable, trait-like features of the human brain, both within and between individuals [1–3]. In particular, ubiquitous patterns of differentiation between sensorimotor and association regions across cortical structure and function seem to represent a major principle of brain organization, also known as the sensorimotor-association (S-A) axis [4, 5]. However, beyond the consistency and robustness of functional networks arranged along this axis lies a subtle –yet notable– degree of intra-individual variability [6], suggesting that the S-A axis also has dynamic properties, even at rest. Given that the brain is an endocrine organ, susceptible to transient endogenous fluctuations in the levels of different steroid hormones across sexes [7], such fluctuations may influence the dynamic reconfiguration of functional networks underpinning intra-individual variability, ultimately supporting flexible cognition and behavior [8]. Neuroendocrine processes are thus likely involved in variability of functional brain organization within an individual in a sex-specific manner [9] – yet how remains unclear.

In the adult mammalian endocrine system, the production of gonadal steroid hormones differs between the sexes. In females of reproductive age, a major source of daily variability in gonadal steroids is dictated by the ovarian cycle, which is responsible for the cyclical production of estradiol and progesterone over the 4-5 day rodent estrous cycle and the monthly human menstrual cycle [10]. In humans both sexes are also subject to cyclical changes in endogenous steroid hormone levels following the 24-hour circadian rhythm, whereby testosterone and cortisol production peaks in the morning and steadily declines throughout the day [11, 12]. Although steroid hormones are not exclusive to either sex, females generally present higher concentrations of estrogens and progesterone, and males generally present higher concentrations of testosterone [13], which explains why research primarily focuses on the predominant hormones of each sex accordingly. Despite these substantial differences in steroid hormone concentrations between males and females, we lack a formal understanding of how sex-specific neuroendocrine mechanisms may interact with human brain organization.

Cross-species evidence points to steroid hormones as potent neuromodulators. Receptors for steroid hormones are expressed throughout the brain, particularly in the hippocampus and medial temporal lobe [14–16]. A foundational study in female rats detected a 30% increase in dendritic spine density of hippocampal CA1 pyramidal neurons on the day of ovulation (when estradiol levels peak) relative to 24 hours later [17], suggesting estradiol’s role in enhancing synaptic plasticity in CA1 neurons [18–21], whilst progesterone appears to inhibit this effect [22]. Androgens, such as testosterone, also appear to influence medial temporal lobe morphology, for example by inhibiting apoptosis in hippocampal neurons [23]. Similarly, findings in humans have linked gonadal steroid levels to changes in brain structure, for example, through effects of estradiol and progesterone levels on hippocampal morphology over the menstrual cycle [24, 25] as well as associations between testosterone levels and cortical thickness during puberty in regions with high androgen receptor density [26]. Moreover, functional magnetic resonance imaging (fMRI) studies have revealed associations between human steroid hormone levels and functional brain activity at rest. Different samples varying in size and sampling frequency suggest that changes in functional connectivity in women are associated with fluctuating levels of endogenous steroid hormones such as estradiol and progesterone over the menstrual cycle [27–29], as well as contraceptive-dependent levels of exogenous steroid hormones [30, 31]. In men, group analyses have revealed changes in resting-state network connectivity related to exogenous increases in testosterone levels [32, 33]. Although considerable evidence from animal and human research supports the role of gonadal steroid hormones in modulating brain structure and function, whether and how sex-specific endogenous fluctuations in steroid hormones contribute to daily variability in functional brain organization remains poorly understood.

Gonadal hormones further have the ability to modulate the stress response through tight interactions between the hypothalamic–pituitary–gonadal (HPG) and hypothalamic– pituitary–adrenal (HPA) axes, which are the neuroendocrine axes respectively producing gonadal and adrenal (i.e., cortisol) hormones [34]. As such, gonadal steroids are thought to contribute to sex differences in the stress response through their activational and organizational effects on the brain throughout the lifespan [35]. For example, circulating estradiol levels in female rodents appear to elevate cortisol levels during both threatening and non-threatening situations, leading to a more robust HPA axis response relative to males [36]. In humans, estradiol levels have also been shown to modulate healthy female functional activity across key regions of the stress circuitry, including the hippocampus, bilateral amygdala, and hypothalamus – an effect that was not observed in women with major depressive disorder, suggesting an association between affective dysfunction and the dysregulation of hormonal effects on stress-related activity [37]. In fact, given that stress contributes to mechanisms of plasticity and vulnerability by physiologically remodeling neural architecture [38], sex differences in the stress response are thought to contribute to differences in the prevalence of affective psychiatric disorders [36]. Moreover, cortisol responsivity seems to both vary [39] and differentially interact with perceived stress [40] at different stages of the menstrual cycle, highlighting the importance of also considering effects related to subjective self-reported cognitive experience. As such, psychosocial and physiological stress levels should be considered as potential neurocognitive and neuroendocrine factors affecting dynamic changes in functional brain organization via sex-specific mechanisms.

In the last decade, dense-sampling –the deep phenotyping of individuals at a high temporal frequency– has emerged as a method to investigate the stability and variability of functional brain organization. Based on the premise that not enough neuroimaging data is collected per individual –yielding estimates with high measurement error [41]– recent initiatives such as the MyConnectome Project [42, 43] and the Midnight Scan Club [44] have demonstrated the utility of dense-sampling. These studies revealed fine-grained features unique to the individual, adding a layer of detail and specificity that is otherwise overlooked in group-averaged data [43]. Aiming to demonstrate the reliability of resting-state functional connectivity patterns, these pioneering studies focused on assessing the within-subject stability –rather than variability– of the functional connectome [2, 45]. As such, they did not investigate the factors and mechanisms that may contribute to intra-individual variability in functional brain activity, nor did they investigate the effects of sex as a biological variable in their analyses [9].

Recently, a dense-sampling approach has been applied on a 23-year-old female (28&Me study; [46]) and a 26-year-old male (28&He study; [47]), who were respectively tested for 30 and 20 consecutive days in time-locked study sessions including brain imaging, venipuncture and salivary sampling, and self-report mood questionnaires. These studies –as well as subsequent studies using the female dataset (e.g., [48–51])– measured day-to-day changes in functional brain activity, reporting associations between hormonal fluctuations and the reorganization of functional networks. However, none of these studies have directly compared intra-individual variability across sexes, nor have they probed and compared sex-specific neuroendocrine effects in relation to major principles of brain organization, such as the S-A axis. In fact, increasing evidence supports the promise of using low dimensional measures of functional connectivity to study variations in sensory-to-association hierarchical patterns of intrinsic cortical organization [5, 52–54]. Conceptually, the S-A axis has been shown to reflect both developmental [4] and evolutionary [55, 56] mechanisms, aligning with microstructural variation [55, 57–59], as well as capturing organizational differences between the sexes [60]. Methodologically, the S-A axis has demonstrated suitable levels of reproducibility, predictive validity, and test-retest reliability [61, 62]. As such, studying daily intra-individual variability along the S-A axis as well as associated sex-specific neuroendocrine factors would allow to contextualize subtle intra-individual changes in the functional connectome at a meaningful organizational level.

In the current work, we capitalize on a dense sampling approach to investigate intra-individual variability along the S-A axis in two healthy young adults, one male and one female, from the aforementioned openly-available datasets [46, 47]), probing the sex-specificity of neuroendocrine factors (i.e., steroid hormone levels), as well as perceived stress, associated with daily variability in functional brain organization. We first applied a dimensionality reduction algorithm to daily functional connectivity matrices in order to compute the S-A axis. After quantifying intra-individual variability along the S-A axis, we directly compared patterns of variability between the female and male participants, and further decoded these patterns with publicly available multimodal brain maps. Next, we probed local and system-level effects of day-to-day changes in hormone levels and perceived stress on the S-A axis in both participants. To do this, we conducted two sets of analyses. First, we specifically assessed effects of steroid hormones that are most predominant within each sex (i.e., estradiol and progesterone in the female participant, testosterone in the male participant), as well as cortisol in the male participant given its availability and given that its production follows circadian fluctuation patterns similar to testosterone. Second, we tested for sex-specific effects of common steroid hormones (i.e., estradiol and testosterone), allowing a direct comparison of effects across the female and male participants. As such, rather than systematically testing for statistical differences between the sexes, our work investigates sex-specific factors that may underpin intra-individual variability along a major principle of functional cortical organization in female and male single individuals.

## RESULTS

### Daily variability in steroid hormone levels and perceived stress

In the female participant (*n* = 29; we excluded experimental day 26 from all analyses given that the corresponding fMRI data collected on that day appeared to be compromised – as further reported in our Methods data inclusion criteria), daily serum steroid hormone fluctuations followed expected patterns throughout the menstrual cycle (Figure 1A). Estradiol levels (mean = 84.26 ± 53.74 pg/mL) showed typical increases and decreases, peaking on day 13 of the menstrual cycle (corresponding to the ovulatory window), whilst progesterone levels (mean = 5.25 ± 5.84 ng/mL) were low during the follicular phase (before ovulation) and high during the luteal phase (after ovulation). In the male participant (*n* = 20), daily salivary steroid hormones showed fluctuating levels of testosterone (mean = 101.61 ± 9.98 pg/mL) and cortisol (mean = 0.50 ± 0.13 ug/dL), which did not follow any expected or cyclical pattern (Figure 1B). Evening experimental sessions allowed to confirm normative circadian patterns of higher testosterone and cortisol levels in the morning relative to the evening in the male participant, although we excluded data acquired during evening sessions to control for time of day in our analyses (see Methods for more detail on our data inclusion criteria).

**Figure 1.**
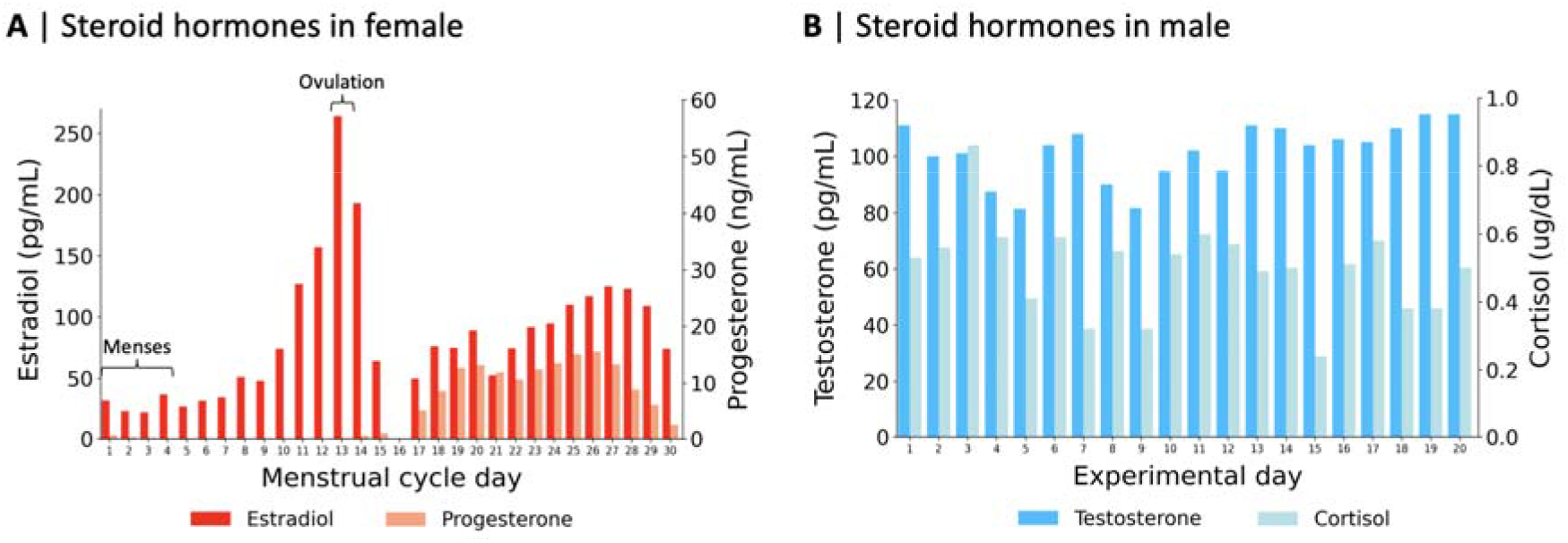
Daily variability in steroid hormone levels. **A** | Estradiol and progesterone serum levels in the female participant across her menstrual cycle (*n* = 29). Note that the menstrual cycle days shown here do not correspond to the experimental sessions, which were rearranged to begin at menstruation for this visualization. Experimental day 26 was removed from this visualization and from all analyses, see Methods for more detail on our data inclusion criteria; **B** | Testosterone and cortisol salivary levels in the male participant across experimental days (*n* = 20).

Self-reported perceived stress was measured with the perceived stress scale (PSS), where PSS scores can range from 0 (low stress) to 40 (high stress). We found no statistically significant difference in PSS scores between the female (mean score = 8.28 ± 6.59) and the male (mean score = 10.10 ± 2.10) participants, as measured by the Mann-Whitney U Test, *U* = 211.5, *p* = 0.11.

### Intra and inter-individual daily variability in functional cortical organization

We computed the S-A axis as our measure of functional cortical organization at each study session in both subjects. For this, we used diffusion map embedding, a non-linear data reduction algorithm, to reduce the dimensionality of the 400x400 functional connectivity matrices, representing the pairwise strength of functional connectivity between Schaefer 400 cortical regions [63]. We thus computed, for each study session within both subjects, the well-replicated principal gradient explaining the most variance in the data (33.95% in the female participant, 35.47% in the male participant) –spanning from unimodal sensorimotor regions to transmodal association regions [5]– which we defined as the S-A axis. Figures 2A and 2B show the mean S-A axes of the female and male participants respectively, computed by applying diffusion map embedding to the mean daily functional connectivity matrices (averaged across study sessions) within each participant. We used the S-A axis to represent functional cortical organization throughout our analyses, where S-A axis loadings represent each of 400 cortical regions’ positions along this low dimensional axis of functional cortical organization.

**Figure 2.**
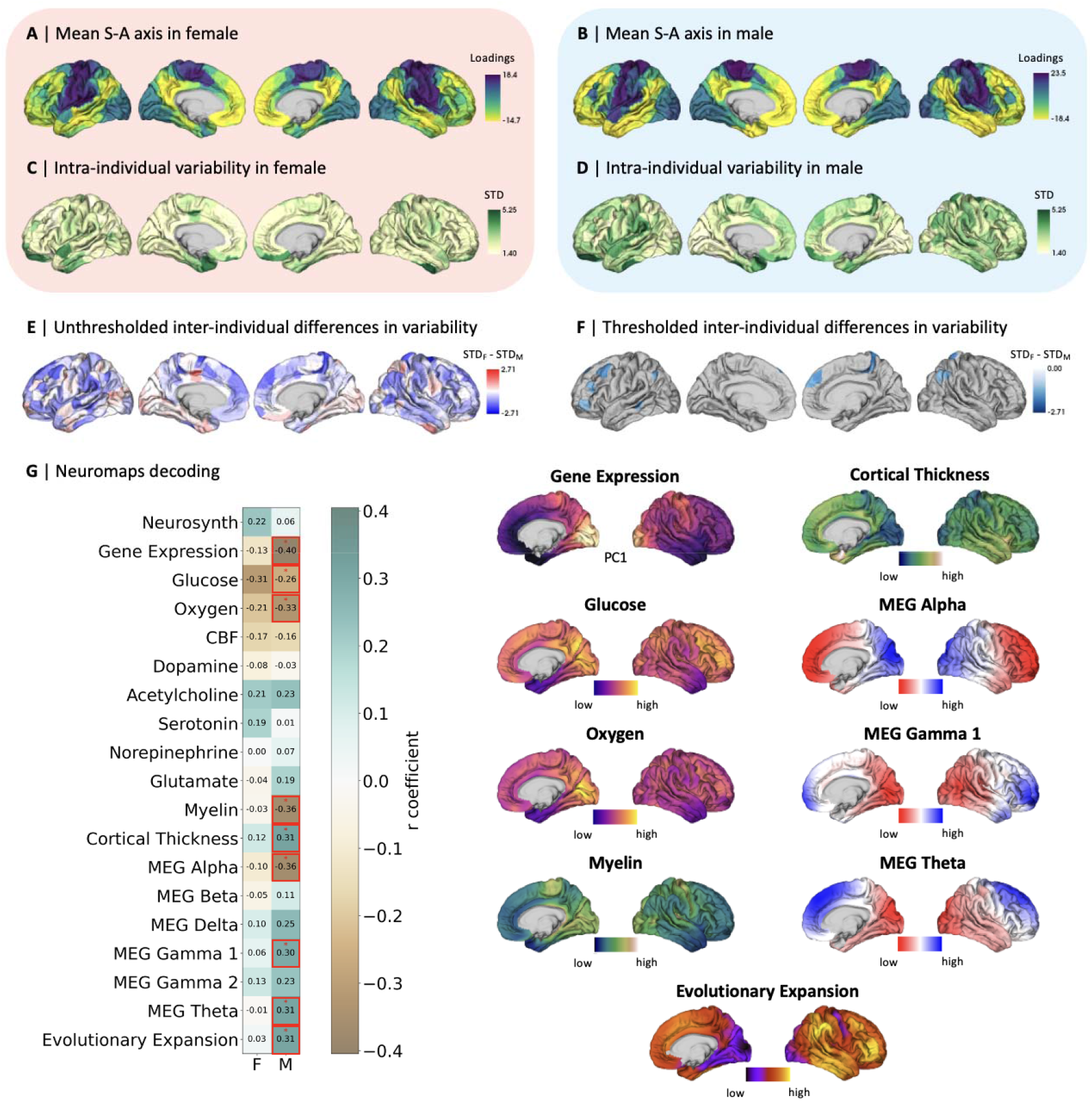
Intra and inter-individual daily variability in functional cortical organization. **A** | Mean sensorimotor-association (S-A) axis loadings across 29 days in the female participant; **B** | Mean S-A axis loadings across 20 days in the male participant; **C** | Intra-individual variability in S-A axis loadings quantified by standard deviation (STD) in the female participant; **D** | Intra-individual variability in S-A axis loadings quantified by STD in the male participant; **E** | Inter-individual differences in intra-individual variability quantified by the subtraction of male from female intra-individual variability; **F** | Thresholded inter-individual differences in intra-individual variability, displaying inter-individual difference in intra-individual variability in false discovery rate (FDR)-corrected parcels (*q* < 0.05) showing statistically significant differences as resulted by the Levene’s test for equality of variances; **G** | Spearman-rank correlations between patterns of intra-individual variability in the female (F) and male (M) participants and 19 brain feature maps sourced from the Neuromaps database, where red * indicates statistically significant correlations after spin permutation testing and FDR correction (*q* < 0.05). Brain feature maps showing statistically significant associations with the male participant’s intra-individual variability are displayed. MEG, magnetoencephalography.

We then probed subtle daily changes in functional cortical organization. For both participants, intra-individual daily variability in S-A axis loadings was quantified using standard deviation (Figures 2C and 2D). We found a statistically significant spatial association between the cortex-wide patterns of female and male intra-individual variability (Spearman’s rank: *r* = 0.29, *p*_spin_ < .001). Both participants displayed greatest variability in temporal limbic and ventral prefrontal regions, extending further across the cortex in the male participant. To quantify inter-individual differences in intra-individual variability, we subtracted male from female standard deviations by cortical parcel (Figure 2E). We then assessed the statistical significance of the inter-individual differences across cortical regions with Levene’s test for equality of variances and found statistically significant greater local intra-individual variability exclusively in the male participant, namely in about 5% of regions (19 out of 400) distributed across functional networks (Figure 2F).

To further interpret intra-individual functional variability, we explored its association to brain features such as gene expression, meta-analytic functional activations subserving behavior and cognition, metabolism, neurotransmitter receptor distribution, brain structure and function (electromagnetic waves), as well as patterns of evolutionary expansion. We thus decoded patterns of intra-individual variability in S-A axis loadings by testing their associations with 19 independent maps of brain features from the publicly available Neuromaps database [64] – see Methods for more information about each map. For this, we computed the Spearman-rank correlation between each brain feature map and both the female and male intra-individual variability maps separately (Figure 2G). Here, we only found statistically significant associations that survived spin permutation testing as well as false discovery rate correction (FDR; *q* < 0.05) for the male participant. Specifically, patterns of male intra-individual variability were negatively associated with patterns of overall gene expression, glucose and oxygen metabolism, myelin, and magnetoencephalography (MEG) alpha activity, illustrating that regions with higher variability are regions typically displaying lower metabolism, myelin intensity, and MEG alpha activity. To note, the directionality of the association with gene expression patterns is irrelevant given that the gene expression map represents the first principal component computed over the expression of 1000 genes from the Allen Human Brain Atlas [65], which should be understood as an axis of variability in the similarity of gene expression profiles rather than a measure with meaningful directionality. Furthermore, patterns of male intra-individual variability were positively associated with patterns of cortical thickness, MEG gamma 1 activity, MEG theta activity, and evolutionary expansion, illustrating that regions with higher variability are regions typically displaying greater cortical thickness, MEG gamma 1 and theta activity, and greater relative cortical surface area expansion from macaque to human.

### Effects of sex-predominant steroid hormones and perceived stress on functional cortical organization

In order to investigate factors potentially underlying dynamic intra-individual daily changes in functional organization, we tested for local effects of steroid hormone levels and perceived stress on the S-A axis loadings by independently fitting different linear models in both participants. Our first set of analyses included steroid hormones that are most predominant within each sex (i.e., estradiol and progesterone in the female participant, testosterone in the male participant), as well as cortisol in the male participant given its availability and given that its production follows circadian fluctuation patterns similar to testosterone. We thus included serum estradiol and progesterone levels as covariates in the linear model for the female participant (*n* = 29), and included salivary testosterone and cortisol levels as covariates in the linear model for the male participant (*n* = 20). Separate additional linear models were used to account for the local effects of perceived stress (PSS score) in both participants independently. Other than 13.5% of cortical regions (54 out of 400) showing statistically significant effects of testosterone on S-A axis loadings in the male participant that were spread across functional networks, no other hormone or stress measure showed local-level effects that survived FDR correction. *t*-maps of all tested local effects are displayed in the Supplementary Figure 1. Furthermore, the female and male participants showed overall different patterns of PSS score effects on S-A axis loadings, as indicated by the lack of association in the spatial patterns of their PSS scores effects on S-A axis loadings, *r* = -0.08, *p*_spin_ = 0.078.

For a more interpretable characterization of daily changes in S-A axis loadings (i.e., local shifts in the position of parcels on the S-A axis), we tested for system-level effects of hormone levels and perceived stress on changes in network topology, that is, changes in the topographical organization of functional networks along the S-A axis (Figure 3A). For this, we independently computed measures of both within-and between-network dispersion across study sessions for both participants as done in previous work [60, 66], based on the Yeo-Krienen seven functional network solution [67]. Within-network dispersion quantifies the spread of cortical regions within each of the seven networks along the S-A axis, with higher values of within-network dispersion indicating higher segregation of regions within a given network. Between-network dispersion quantifies the pairwise distance between a given pair of functional networks along the S-A axis, with higher values of between-network dispersion indicating a higher segregation of the two given networks. We thus computed measures of within-network dispersion for all seven functional networks, and measures of between-network dispersion for all possible pairwise combinations of the seven networks (i.e., 21 pairs of networks in total). Then, we fitted the same linear models used to test for local effects on the S-A axis, separately testing for steroid hormone and perceived stress effects on all measures of within-and between-network dispersion. Overall, we found two statistically significant effects: An association between testosterone and segregation within the dorsal attention network in the male participant, *t* = 2.943, *p*_spin_ = 0.005, as well as an association between progesterone and integration between the frontoparietal and default mode networks in the female participant, *t* = -3.792, *p*_spin_ = 0.008. In Figure 3B-E, we summarized patterns of all tested system-level effects on functional network dispersion along the S-A axis, using heatmaps to highlight the directionality of effects, where positive *t*-values are illustrated in purple and negative *t*-values in brown, respectively indicating segregation and integration effects. Broadly, in the female participant, estradiol displayed patterns of greater segregation within sensory networks and greater integration within association networks, whereas progesterone displayed almost exclusively patterns of greater integration within networks (only displaying associations with greater segregation in the dorsal attention network). In the male participant, testosterone almost exclusively displayed patterns of greater segregation within networks, which also translated to overall greater segregation between networks, whilst cortisol was mainly associated with greater within-network integration. Effects of perceived stress tended towards integration both within and between functional networks, with more pronounced effects in the female participant relative to the male. The detailed statistical results for all analyses of system-level effects on functional organization are summarized in Supplementary Tables 1-3.

**Figure 3.**
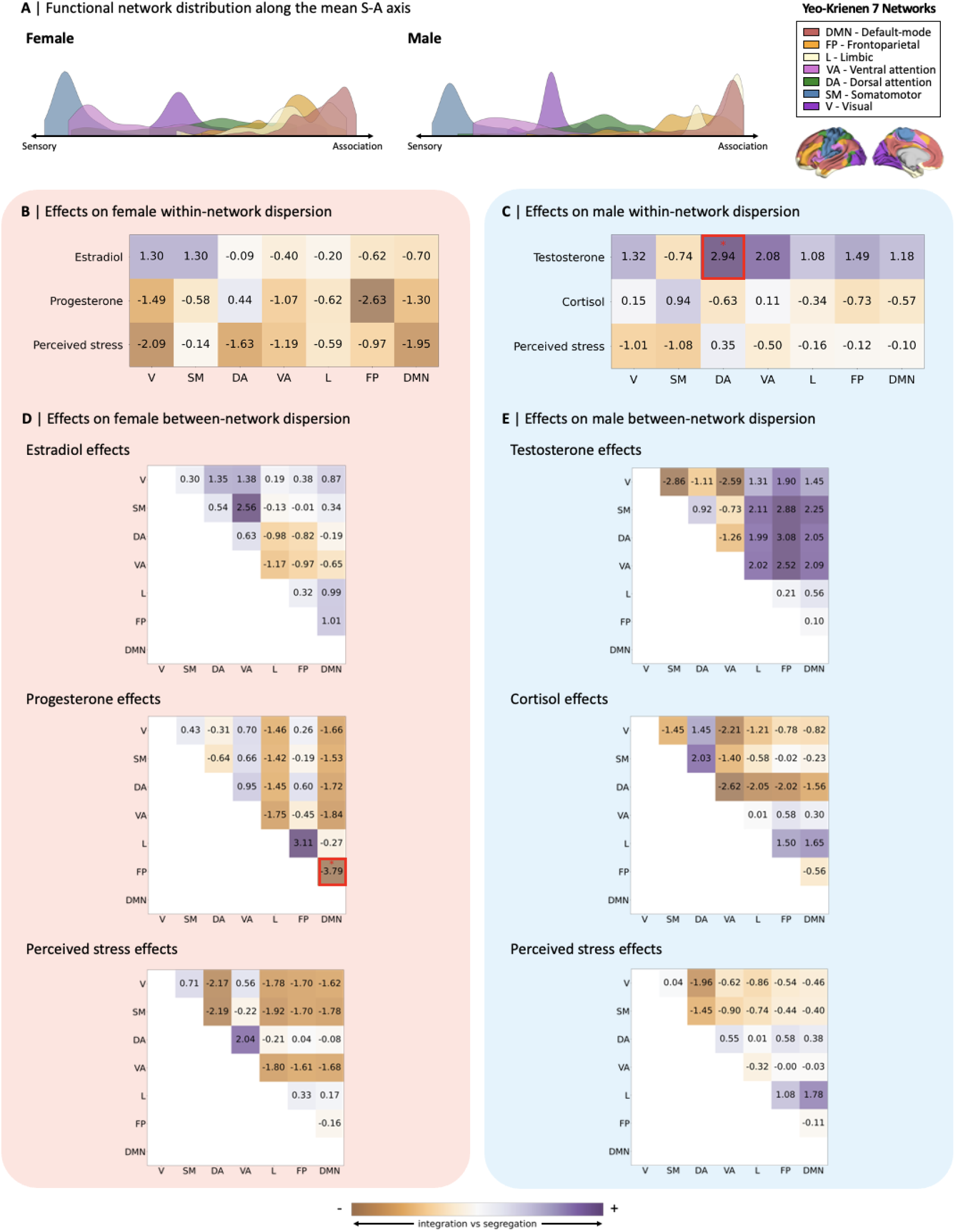
System-level effects of sex-specific steroid hormones and perceived stress on functional cortical organization. **A** | Visualization of the distribution of the seven Yeo-Krienen functional networks along the mean sensorimotor-association (S-A) axis in the female and male participants. Heatmaps summarizing the *t*-values for system-level effects across functional networks of estradiol, progesterone, and perceived stress on the female participant’s within-(**B**) and between-(**D**) network dispersion, and effects of testosterone, cortisol, and perceived stress on male participant’s within-(**C**) and between-(**E**) network dispersion. *t*-values were obtained from linear models including different sets of covariates, namely estradiol and progesterone (for female hormone effects), testosterone and cortisol (for male hormone effects), and perceived stress only (for both female and male, tested separately). Red * indicates statistical significance of effects corrected for multiple comparisons, at Bonferroni-corrected thresholds of *p* < 0.004 (0.025/7) for the within-network effects and *p* < 0.001 (0.025/21) for the between-network effects, as well as corrected for spatial autocorrelation via spin-permutation testing (1000 permutations). Positive *t*-values represent higher segregation and negative *t*-values represent higher integration effects. V, visual; SM, somatomotor; DA, dorsal attention; VA, ventral attention; L, limbic; FP, frontoparietal; DMN, default-mode network.

### Sex-specific effects of common steroid hormones on functional cortical organization

After testing for the hypothesized effects of steroid hormones that are most predominant within each sex, our second set of analyses tested for sex-specific effects of common steroid hormones (i.e., estradiol and testosterone) on functional organization in both participants, to allow a direct comparison of effects across participants. We thus included estradiol and testosterone as covariates in a new model that we independently tested for each participant in order to directly compare the sex-specific local-level effects of these common steroid hormones on S-A axis loadings. To increase comparability between participants, we used serum hormone levels for the male participant in this set of analyses, at the cost of a smaller sample size (*n* = 15; see Methods section for more detail on our data inclusion criteria and the availability of steroid hormones per subject). Similar to the models testing for sex-specific steroid hormone effects, we did not find any statistically-significant local-level effects following our statistical corrections. We compared brain-wide patterns of local effects (i.e., unthresholded *t*-values) of estradiol in the female (Figure 4A) and male (Figure 4B) participants, which revealed no statistically significant association (Figure 4C; *r* = -0.15, *p*_spin_ = 0.11). The comparison of brain-wide patterns of local effects of testosterone in the female (Figure 4D) and male (Figure 4E) participants revealed a small statistically significant negative association (Figure 4F; *r* = -0.23*, p*_spin_ = 0.04). Both of these correlation findings suggest differences in the female and male patterns of local-level effects of estradiol and testosterone on functional organization.

**Figure 4.**
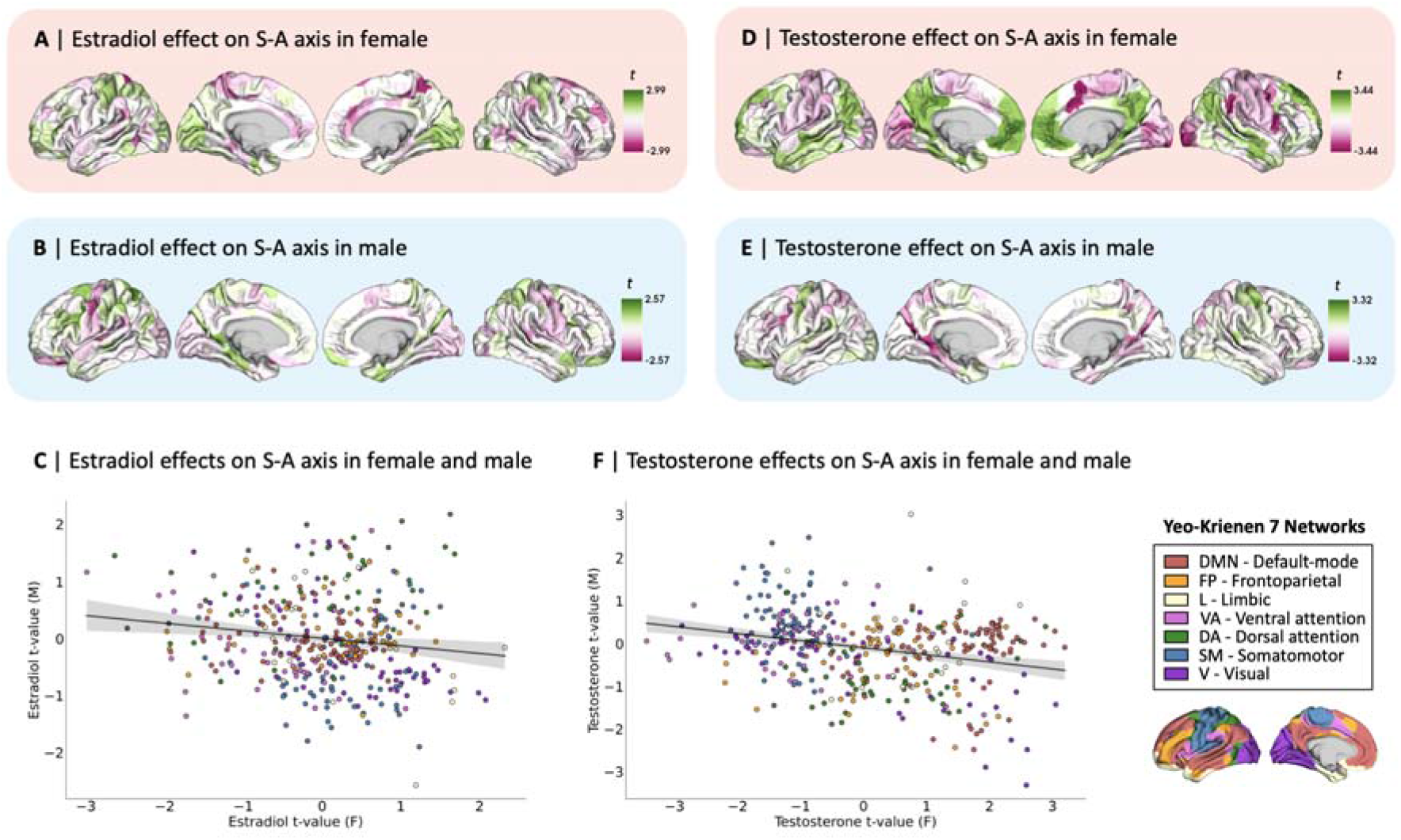
Local-level effects of estradiol and testosterone on functional organization in female and male participants. For both sexes, effects were tested in linear models including estradiol and testosterone as covariates. Unthresholded *t*-maps of linear regression results showing patterns of local effects of estradiol on sensorimotor-association (S-A) axis loadings in the **A** | female and **B** | male participants. **C** | Scatterplot displaying the spatial correlation between patterns of local estradiol effects on S-A axis loadings in female participant (F; x-axis) and in male participant (M; y-axis), *r* = -0.15, *p*_spin_ = 0.11; colors denote the seven Yeo-Krienen functional networks. Unthresholded *t*-maps of linear regression results showing patterns of local effects of testosterone on S-A axis loadings in the **D** | female and **E** | male participants. **F** | Scatterplot displaying the spatial correlation between patterns of local testosterone effects on S-A axis loadings in female participant (x-axis) and in male participant (y-axis), *r* = -0.23, *p*_spin_ = 0.04.

Again, for a more interpretable characterization of daily changes in S-A axis loadings, we tested for system-level effects of estradiol and testosterone on changes in the topographical organization of functional networks along the S-A axis (Figure 5A), and did not find any statistically significant effects of steroid hormones on within or between-network dispersion. In Figure 5B-E, we summarized patterns of all system-level effects on functional network dispersion along the S-A axis, using heatmaps to highlight the directionality of effects. Here, we observed distinctly diverging patterns effects between the two subjects. For example, estradiol in the male participant showed greater integration within sensory networks and greater segregation within association networks, opposite to female patterns as previously described. The detailed statistical results for system-level effects of estradiol and testosterone on functional organization in both participants are summarized in Supplementary Tables 4 and 5.

**Figure 5.**
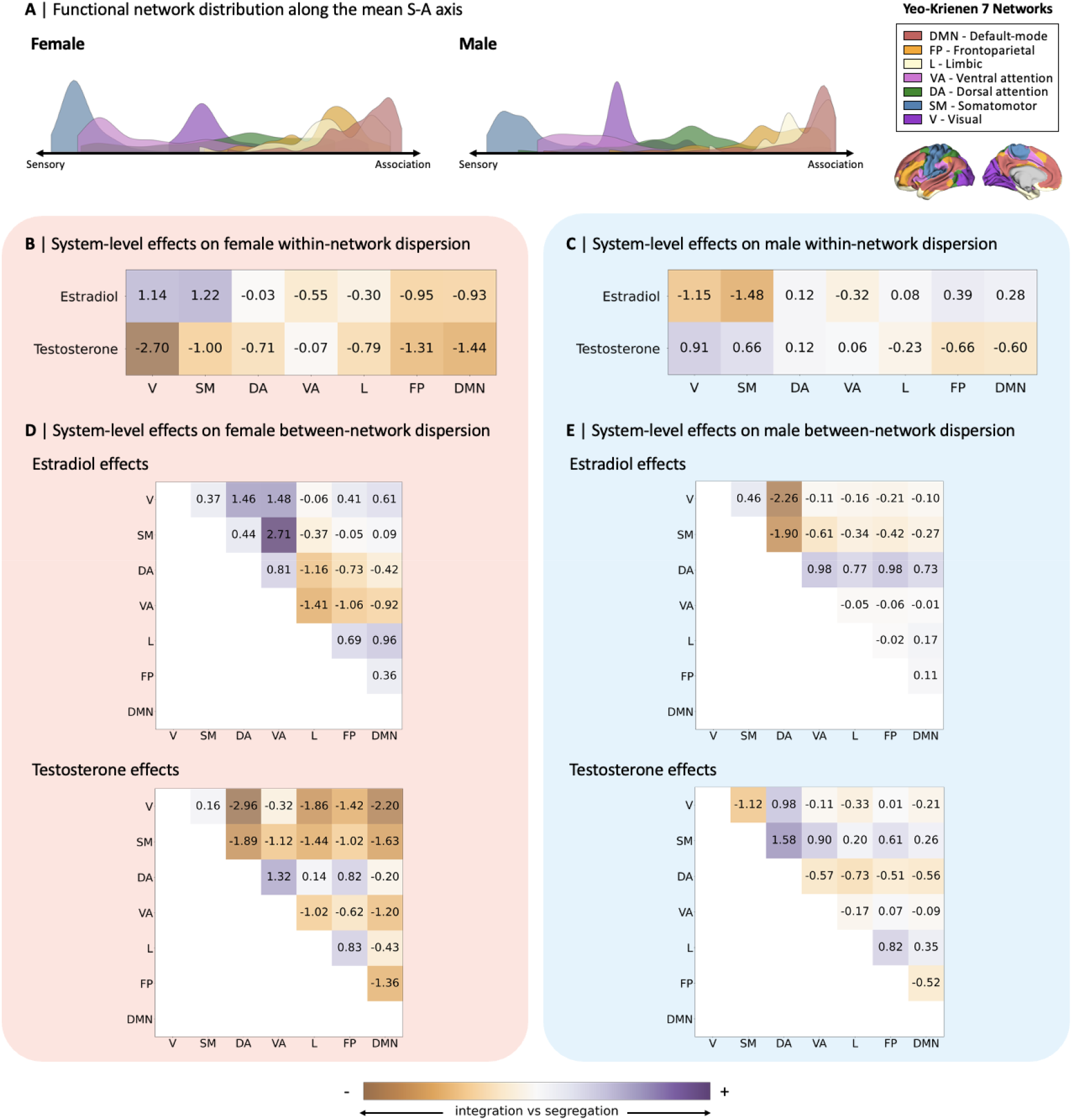
System-level effects of estradiol and testosterone on within-and between-network dispersion in female and male participants. **A** | Visualization of the distribution of the seven Yeo-Krienen functional networks along the mean sensorimotor-association (S-A) axis in the female and male participants. Heatmaps summarizing the *t*-values for system-level effects across functional networks of estradiol and testosterone for the female participant’s within-(**B**) and between-(**D**) network dispersion, and the male participant’s within-(**C**) and between-(**E**) network dispersion. *t*-values were obtained from linear models including estradiol and testosterone as covariates in both the female and male models, tested separately per participant. None of the tested effects were statistically significant after correction for multiple comparisons, i.e., at Bonferroni-corrected thresholds of *p* < 0.004 (0.025/7) for the within-network effects and *p* < 0.001 (0.025/21) for the between network effects. Positive *t*-values represent higher segregation and negative *t*-values represent higher integration effects. V, visual; SM, somatomotor; DA, dorsal attention; VA, ventral attention; L, limbic; FP, frontoparietal; DMN, default-mode network.

## DISCUSSION

In the current work, we used a dense-sampling approach to investigate sex-shared and sex-specific neuroendocrine factors that may be associated with intra-individual daily variability in functional brain organization, directly comparing variability and associated effects across two deeply phenotyped young adults, one female and one male. Different from previous work using dense-sampling, we computed a low dimensional representation of patterns of resting-state functional connectivity –the S-A axis, spanning from unimodal sensorimotor regions to transmodal association regions– to quantify subtle daily intra-individual variability along this key hierarchical principle of functional cortical organization. Overall, participants showed unique cortical patterns of intra-individual variability in S-A axis loadings, with similar cortical areas (i.e., temporal limbic and ventral prefrontal regions) displaying the largest amount of variability across participants and male variability extending further across the cortex. We also found statistically significant greater intra-individual variability exclusively in the male relative to the female participant, as well as associations between male whole-brain patterns of intra-individual variability and a range of brain features including brain metabolism, structure, electrophysiology, genetics, and phylogeny. When testing for local-and system-level effects of steroid hormones and perceived stress on functional organization, we found only a few effects that survived our statistical corrections, which we suspect to be a consequence of low statistical power. However, we observed distinctly diverging patterns of hormone and perceived stress effects on network topology in the female and male participants under study. Collectively, our findings suggest subtle inter-individual differences in intra-individual daily variability along a major principle of functional cortical organization and hint at potential sex-specific neuroendocrine processes that require testing in larger samples.

By establishing daily intra-individual variability in the functional cortical organization, steroid hormone levels, and perceived stress of two densely sampled individuals [46, 47], our findings highlight the dynamic nature of brain function, embedded in equally dynamic endocrine and cognitive systems (i.e., the steroid hormone levels and perceived stress under study). We found that patterns of intra-individual variability did not follow a particular sensory to association differentiation, unlike previous reports of greater within-subject variability in lower order unimodal regions and greater between-subject variability in higher order transmodal regions [42, 68]. Nevertheless, intra-individual variability was greatest in temporal limbic and ventral prefrontal regions in both participants, which to some extent replicates previous findings of greater variability in the limbic network of 30 densely-sampled individuals [69]. Although greater variability in the limbic network may in part reflect the lower signal-to-noise ratio typically observed in temporal regions during fMRI scans [70], limbic regions are also known for their remarkable plasticity, which has been linked to laminar patterns of structural variability [71]. The temporal lobe is a cortical area that is particularly dense with steroid receptors [14–16], whose volume has been shown to vary as a function of steroid hormone levels [24, 25, 72, 73]. Altogether, our findings suggest unique whole-brain patterns of subtle daily intra-individual changes along a major principle of functional cortical organization, with some similarities across participants in the regions displaying the greatest amount of variability.

Higher variability in S-A axis loadings was exclusively observed in the male participant when statistically testing for inter-individual differences in intra-individual variability along this low dimensional measure of functional cortical organization. Given that we only sampled one individual of each sex, we cannot generalize this finding to the group level. Nevertheless, an evolutionary hypothesis supporting greater male variability in both biological and cognitive phenotypes has long been formulated [74] and has more recently been empirically supported by different measures of brain structure across the lifespan [75–77]. Greater male variability is thought to potentially result from constraints imposed by genetic architecture, namely the heterogametic nature of male sex chromosomes (XY) as opposed to identical sex chromosomes in females (XX) [78]. In fact, our exploratory analyses showed that intra-individual variability in the male participant was further associated with patterns of overall gene expression, as well as patterns of glucose and oxygen metabolism, myelin, cortical thickness, MEG alpha, MEG gamma 1, MEG theta activity, and evolutionary expansion. It is important to note that these different brain features were obtained from openly available datasets representing group averages rather than being specific to the individuals under study. Yet, these multilevel features are theoretically pertinent to functional organization and may thus plausibly contribute to intra-individual functional variability, as suggested by our findings. For example, metabolic substrates such as glucose and oxygen are directly related to the brain’s energy expenditure and local changes in hemodynamics, thus relevant to the measurement of the blood-oxygen-level-dependent (BOLD) signal [79]. Furthermore, the comparison of MEG and fMRI signals –i.e., local field potentials and the BOLD response, respectively– is conceptually plausible given that both predominantly pertain to post-synaptic (dendritic, rather than axonal) signalling [80]. Finally, patterns of evolutionary cortical expansion have previously been shown to reflect spatial patterns of variability along the S-A axis [55, 56, 81] as well as inter-individual variability in functional connectivity [68]. Interestingly, the fact that associations between intra-individual variability and the tested brain features were only statistically significant in the male participant suggests that sources of variability may differ between the two participants. We indeed only observed a minor association in patterns of intra-individual variability between our participants, consistent with previous research suggesting that –beyond some shared patterns of variability– a larger proportion of intra-individual daily changes in functional organization is unique to the individual [42, 68]. Our findings thus point to unique multilevel factors associated to intra-individual variability –possibly in a sex specific manner– which should be further investigated in larger samples.

To probe the possible multilevel sex-specific underpinnings of daily variability along our low dimensional measure of functional cortical organization, we tested both local-and system-level effects of steroid hormone levels and self-reported perceived stress in the female and male participants separately. Overall, we found a few effects that survived our statistical corrections. At the local level, we observed effects of testosterone on S-A axis loadings in the male participant, which were spread across functional networks. At the system level, we observed effects of testosterone on male dispersion within the dorsal attention network (where testosterone levels were associated with increased segregation, indicating an increase in network spread along the S-A axis), as well as effects of progesterone on female dispersion between the frontoparietal and default mode networks (where progesterone levels were associated with increased between-network integration, indicating an increase in similarity of these functional communities on the S-A axis). We suspect that the scarcity of statistically significant results may altogether be attributed to low power, given that samples ranged from 15 to 29 datapoints across analyses. Nonetheless, we observed strikingly diverging patterns of steroid hormone effects on network topology between the female and male participants (i.e., hormones showing distinct associations with network integration versus segregation). In fact, by using a low dimensional representation of functional organization and measuring changes in functional network topology with a measure of network dispersion, we can meaningfully understand and interpret the topological changes as functional communities becoming more similar or different along the hierarchical domains of the S-A axis [60, 66]. Functional network integration and segregation are more generally considered to be important indicators of network topological structure and reconfiguration underpinning cognition, as they have been associated with a range of cognitive functions as well as changes in brain states, arousal, and energy expenditure [8]. Moreover, even more intriguing were the distinctly opposite patterns of effects of common hormones tested in the female and the male participants. Whole-brain spatial patterns of local effects were either statistically uncorrelated (i.e., effects of estradiol) or negatively correlated (i.e., effects of testosterone) between participants. System-level effects of estradiol on within-network dispersion also showed opposite patterns between participants, that is, greater segregation within sensory networks and greater integration within association networks in the female participant, as opposed to the male participant who showed the opposite patterns. Similarly, patterns of local perceived stress effects on the S-A axis loadings were uncorrelated between the participants, and the qualitative comparison of perceived stress system-level dispersion effects across participants further indicates more pronounced effects in the female participant relative to the male. This suggests the possibility of different stress mechanisms between the participants, possibly in line with previous findings of varying psychophysiological effects of cortisol levels and perceived stress throughout the menstrual cycle [39, 40]. Altogether, our findings underscore heterogeneous and widespread patterns of effects that are not specific to a set of regions or networks. The divergence of findings between the participants further suggests individual differences in neuroendocrine and stress effects on network topology, hinting at potentially sex-specific processes that require systematic and statistical testing in larger samples.

Despite the insights gained through our study, some limitations should be acknowledged. First, our small sample of two young healthy adults –one female and one male– not only entails low statistical power, likely underpinning the scarcity of statistically significant effects in our study, but also precludes the generalization of results to the population-level. We used data that was collected with a high sampling frequency, conscious of the trade-off of data depth (i.e., repeated deep phenotyping in the same individuals) over breadth (i.e., across multiple individuals). As previously eloquently formulated: “Just as no single brain is representative of a population, no group-averaged brain represents a given individual” [42]. Our focus was thus not to yield generalizable findings per se, but to probe fine-grained intra-individual effects that provide a multilevel account of factors potentially influencing intra-individual variability in functional organization. In fact, recent work from our group has highlighted sex differences in functional organization in a large sample (*N* = 1000), which could not be explained by differences in cortical morphometry [60], requiring a deeper investigation of other potential explanations, such as neuroendocrine and neurocognitive factors. Although our group has also observed sex differences in isocortex and hippocampus microstructure related to self-reported female menstrual cycle stage and hormonal contraceptive use [82], those measures were only proxies of female hormone levels, and establishing neuroendocrine mechanisms strictly requires endocrine samples. Our current study thus allowed us to bridge both methodological and conceptual gaps with a deeply phenotyped sample, providing insights on intra-individual variability that was not otherwise possible in large samples and thus highlighting the complementary nature of both approaches [41].

Second, with respect to interpreting sex effects in our current study, we cannot determine the extent to which the observed inter-individual differences in intra-individual variability may be explained by sex-specific mechanisms as opposed to broader individual differences. By only including one individual of either sex, we cannot assume the degree to which our participants may be representative of the greater female and male populations respectively, with increasing evidence further suggesting that sex should actually be treated as a continuous variable in biological research [83, 84]. Furthermore, we did not consider possible effects of steroid hormones on functional organization in gender-diverse individuals, who further challenge the notion of binary female-male categories. In fact, steroid hormone levels are not fixed and may be dynamically affected by gendered social experiences [85], as well as a range of more general environmental factors that go beyond sex and gender-identity, such as sleep, nutrition, caffeine consumption, physical exercise, and stress [86–94]. As such, a larger sample capturing greater variability across females, males, and individuals in general is necessary to establish the degree of sex-specificity of the effects tested in our study, as well as to further assess possible gender-specific effects. Our study is however novel in exploring such sex-specific neuroendocrine effects on the S-A axis in densely sampled individuals.

Third, it could be contended that our findings of intra-individual daily variability in functional cortical organization are capturing random noise in our data. However, we explicitly made methodological decisions aimed at reducing biases and noise particularly pertaining to daily variability and endocrine effects. First, we used the S-A axis as our measure of functional cortical organization, which –through its low dimensionality and thresholding– has been shown to have greater test-retest reliability than more commonly used measures of unthresholded edge-wise functional connectivity [61, 62]. In terms of data design, study sessions were time-locked, and food and caffeine intake prior to study sessions was strictly controlled through abstinence in order to limit confounding physiological effects [46, 47]. Steroid hormones are also thought to potentially induce physiological artefacts, such as local changes in cerebral blood perfusion [95], which could be mistaken as cognitively-pertinent changes in brain function [96]. However, various steps were taken in the preprocessing of our fMRI data –such as global signal scaling and linear detrending of voxel-wise timeseries– to account for temporal and spatial fluctuations in signal intensity and to remove the effects of head motion and physiological noise features such as cerebral spinal fluid from the BOLD signal. We also used coherence as our measure of functional connectivity, which is known for its robustness to temporal variability in regional hemodynamics as well as its measurement of timeseries covariances in frequencies outside the spectrum prone to contamination by physiological noise [97]. As such, the intra-individual daily changes in variability observed in our study are likely to reflect meaningful fluctuations in signal beyond noise.

All in all, by observing subtle daily changes along a low dimensional measure of functional cortical organization in two densely sampled healthy young adults, co-occurring with fluctuations in steroid hormone levels and perceived stress, our findings underscore the importance of holistically considering the brain as an organ embedded in an extensive network of interacting endocrine and psychophysiological systems. By observing diverging patterns of effects in a female and a male participant, we highlight the need for research to systematically test for sex effects, particularly considering the sex-specificity of neuroendocrine mechanisms [9]. Importantly, by showing that a male individual is as subject to hormone-related fluctuations in functional brain organization as a female, we debunk the deeply rooted belief that endocrine variability is an exclusively female concern, which has led to the historical underrepresentation of women from research studies [98]. Going forward, giving equal consideration to both sexes –as well as combining dense sampling approaches with large population-based studies– is necessary to gain a thorough understanding of neuroendocrine and neurocognitive processes underlying variability along principles of functional brain organization in health and disease.

## METHODS

The current study relies on the use of open data, whose methods have already been reported elsewhere in detail (see [46, 47] for the original publications).

### Participants and study design

Our sample (*N* = 2) consisted of one female (23 years) and one male (26 years), both right-handed and Caucasian, with no history of endocrine disorders, neuropsychiatric diagnoses, or head injuries. The female participant reported regular menstrual cycles (occurring every 26-28 days, with no missed periods) and refrained from taking hormone-based medication in the 12 months preceding data collection. Participants gave written informed consent for studies that were originally validated by the University of California, Santa Barbara Human Subjects Committee.

The original study designs for the collection of the female and male data slightly differed and are fully reported in [46] and [47], respectively. Here, we report the original and complete study designs although we use only part of the collected data for our analyses in order to maximize consistency and comparability between the participants (see our data inclusion criteria below). For 30 consecutive days, both participants underwent behavioral assessments, assessments for hormone analysis (the female participant underwent exclusively serological assessments, whereas the male participant underwent a combination of serological and salivary assessments), and brain structural and functional magnetic resonance imaging (MRI) in time-locked sessions. Experimental sessions for the female participant occurred exclusively in the morning, whereas sessions for the male participant occurred in the morning for the first 10 days, both in the morning and evening for the following 10 days, and in the evening for the last 10 days, for a total of 40 sessions. Given that blood samples could only be drawn once per day, the male participant’s serological assessments were conducted in the morning session for the first 15 days and in the evening session for the last 15 days, whilst salivary samples were collected at every session. Each session started with a behavioral assessment consisting of self-report questionnaires including the Perceived Stress Scale (PSS; adapted to reflect past 24 hours), consisting of 10 questions measuring the level of appraised stress from life situation on a five point Likert scale from 0 (“never”) to 4 (“very often”), for a total PSS score ranging from 0 (low stress) to 40 (high stress) [99].

The time-locked collection of steroid hormones was subsequently carried out. For the female participant, steroid hormone samples were collected exclusively via venipuncture, beginning at 10:00 a.m. ± 30 minutes, whereas salivary sampling followed by venipuncture was used to collect the male participant’s steroid hormones, beginning at 7 am for morning sessions and at 8 pm for evening sessions. Following safety guidelines, blood was drawn only once on days with two sessions (i.e., in the morning for experimental days 1-15 and in the evening for days 16-20). Endocrine samples were collected after abstaining from food or drink consumption (including caffeine and excluding water) for at least 2h (female participant), at least 8h (male participant, morning sessions), and at least 1.5 hours (male participant, evening sessions).

### Steroid hormone measurements

For the female participant, serum levels of gonadal steroid hormones (17β-estradiol, progesterone, and testosterone), as well as pituitary gonadotropins (luteinizing hormone (LH) and follicle stimulating hormone (FSH)), were sampled. For the male participant, both serum and salivary levels of total testosterone and cortisol were sampled, as well as serum levels of 17β-estradiol. The saliva sample (∼2mL) was collected over 5-10 min of passive drooling at every session, before storing the sample in a plastic cryovial at -20°C until assayed. Saliva concentrations of testosterone and cortisol were determined using enzyme immunoassay at the Brigham and Women’s Hospital Research Assay Core.

For both participants, a 10 cc mL blood sample was collected per session by a licensed phlebotomist via the insertion of a saline-lock intravenous line into the dominant or non-dominant forearm and the use of a vacutainer SST (BD Diagnostic Systems). For both participants, the serum samples were first allowed to clot at room temperature for 45-60 min, then centrifuged (2100 x g for 10 min) and aliquoted into three 2 mL microtubes. The samples were then stored at -20 °C until assayed. At the Brigham and Women’s Hospital Research Assay Core, liquid chromatography-mass spectrometer (LCMS) was used to determine serum concentration for all steroid hormones, and an immunoassay was used to determine serum concentration for all gonadotropins in the female participant (i.e., follicle stimulating hormone (FSH) and luteinizing hormone (LH)).

Assay sensitivities, dynamic range, and intra-assay coefficients of variation respectively were as follows: estradiol, 1 pg/mL, 1–500 pg/mL,[<[5% relative standard deviation (RSD); progesterone, 0.05 ng/mL, 0.05–10 ng/mL, 9.33% RSD; testosterone, 1.0 ng/dL,[<[4% RSD; testosterone, 1.0 ng/dL, 1–200 ng/dL, <2% RSD; cortisol, 0.5 ng/mL, 0.5-250 pg/mL, <8% RSD. Gonadotropin levels were determined using chemiluminescent assay (Beckman Coulter), with assay sensitivity, dynamic range, and the intra-assay coefficient of variation as follows: FSH, 0.2 mIU/mL, 0.2–200 mIU/mL, 3.1–4.3%; LH, 0.2 mIU/mL, 0.2–250 mIU/mL, 4.3–6.4%.

### MRI acquisition

Both participants underwent a 1h-long MRI scan at every session, conducted on a Siemens 3T Prisma scanner with a 64-channel phased-array head coil. Structural anatomical images were acquired using a T1-weighted magnetization prepared rapid gradient echo (MPRAGE) sequence (TR = 2500 ms, TE = 2.31 ms, TI = 934 ms, flip angle = 7°, 0.8 mm thickness) and a gradient echo fieldmap (TR = 758 ms, TE1 = 4.92 ms, TE2 = 7.38 ms, flip angle = 60°). Resting-state fMRI images were acquired with a T2*-weighted multiband echo-planar imaging (EPI) sequence measuring the blood oxygenation level-dependent (BOLD) contrast (TR = 720 ms, TE = 37 ms, flip angle = 52° (female participant) and 56° (male participant), multiband factor = 8; 72 oblique slices, voxel size = 2 mm3). The resting-state scans lasted 10 and 15 min for the female and male subjects respectively. To reduce head motion, both participants’ heads were secured in a 3D-printed custom-fitted foam head case. Overall head motion, characterized by mean framewise displacement (FWD), was minimal for both participants.

### fMRI preprocessing

The preprocessing of fMRI data was performed in MATLAB using the Statistical Parametric Mapping 12 (SPM12, Wellcome Trust Centre for Neuroimaging, London) software and is fully reported by [46] and [47]. The preprocessing pipeline was identical for both participants. In short, to correct for head motion and geometric deformations, functional images were realigned and unwarped, followed by a co-registration of the mean motion-corrected images to the anatomical images. The Advanced Normalization Tool’s (ANTs) multivariate template construction was used to normalize all scans to a subject-specific template [100]. The functional data was subsequently smoothed using a 4 mm full-width half maximum (FWHM) isotropic Gaussian kernel. To account for fluctuations in signal intensity across time and space, global signal scaling (median = 1000) was applied and voxel-wise timeseries were detrended linearly. After removing the effects of five sources of physiological noise (cerebrospinal fluid and white matter signal) as well as head motion, the residual BOLD signal was extracted from each voxel. A Volterra expansion of translational/rotational motion parameters was used to model head motion based on the Friston-24 approach, which accounts for the non-linear and autoregressive effects of head motion on the BOLD signal [101]. In the current study, we did not apply further global signal regression.

### Functional connectivity and the S-A axis of functional organization

Throughout this work, we used the Schaefer 400-region cortical parcellation [63] as well as its associated Yeo-Krienen seven functional network solution including the visual, somatomotor, dorsal attention, ventral attention, limbic, frontoparietal, default-mode networks [67]. As reported by [46] and [47], the first eigenvariate across functional volumes was used to extract a summary time series in order to compute functional connectivity for each scanning session. The spectral association between time series data from each region was estimated with magnitude-squared coherence, yielding a 400x400 functional connectivity matrix for each experimental session indicating the strength of functional connectivity between all pairs of regions (FDR-thresholded at *q* < 0.05). Coherence was chosen to measure interregional functional connectivity because it avoids contamination by physiological noise given that it is not sensitive to the shape of the regional hemodynamic response function, which can vary as a functional of vascular differences [97].

We then applied diffusion map embedding, a non-linear dimensionality reduction algorithm, on the functional connectivity matrices in order to generate low-dimensional representations of macroscale functional organization [5]. Diffusion map embedding compresses high-dimensional data into low-dimensional “gradients” or axes describing the global structure of the data, along which data points that are highly associated are clustered closer together (i.e., they have similar loadings on the axes), and data points that have low association are further apart [102]. To this end, we used the BrainSpace Python toolbox [103] to generate 10 gradients with the following parameters: 90% threshold (i.e., only considering the top 10% row-wise z-values of functional connectivity matrices, representing each seed region’s top 10% of maximally functionally connected regions), α = 0.5 (α controls whether the geometry of the set is reflected in the low-dimensional embedding – i.e., the influence of the sampling points density on the manifold, where α = 0 (maximal influence) and α = 1 (no influence)), and t = 0 (t controls the scale of eigenvalues). First, for both participants separately, mean gradients were computed by reducing the dimensionality of their mean functional connectivity matrices (i.e., averaged across study sessions). Then, using the same parameters, we computed “daily” gradients, i.e., for each scanning session. In order to maintain comparability for intra-individual analyses, the daily gradients were aligned to their respective mean gradients (i.e., per participant) using Procrustes alignment. Finally, for data from each experimental session, we took the well-replicated principal gradient explaining the most variance in the data and spanning from sensory to association regions [5], which we labeled the S-A axis and used to represent functional cortical organization. In our analyses, we refer to S-A axis loadings, which represent each cortical region’s position on the S-A axis.

### Data inclusion

The female subject’s fMRI data collected on experiment day 26 appeared to be compromised and was thus excluded from our analyses, as further noted in [46]. Furthermore, considering that study designs slightly differed for the two participants, we conducted our analyses on only a part of the data that was originally collected. In our first set of analyses, for the female participant, we chose to include serum levels of estradiol and progesterone, as these steroid hormones are the most potent and studied endocrine neuromodulators in females. For the male participant, we chose to include morning salivary levels of testosterone, as this steroid hormone is a more potent endocrine neuromodulator in males, as well as cortisol given its availability and given that its production follows circadian fluctuation patterns similar to testosterone. Furthermore, we chose morning salivary samples for the male participant (rather than serum/evening samples) in this first set of analyses in order to maximize our sample size (*n* = 20 timepoints) whilst maintaining intra-individual consistency and keeping the time of data collection comparable between participants. Although serum hormone measurements are known to be more accurate, we confirmed the validity of the salivary hormone measurement (and thus their comparability to serum levels) in the male participant by correlating serum and salivary levels for testosterone *(r* = 0.90, *p* = 0.001) and cortisol (*r* = 0.92, *p* = 0.001). In our second set of analyses, aimed at comparing the local-and system-level effects of common steroid hormones (i.e., estradiol and testosterone) between participants, we used morning serum hormone levels for the male participant in order to increase comparability with the female subject at the cost of decreasing male sample size (*n* = 15).

### Statistical analyses

For each participant, intra-individual daily variability in functional organization was computed by taking the standard deviation of each parcel’s S-A axis loading across study sessions. Spearman-rank correlation was used to test the similarity of the two participants’ intra-individual variability maps, followed by a spin-permutation test with 1000 spherical rotations to control for spatial autocorrelation [104]. To quantify inter-individual differences in intra-individual variability, we subtracted the standard deviations of male S-A axis loadings from female S-A axis loading within each parcel and then assessed the statistical significance of the inter-individual differences with Levene’s test for equality of variances, on which we further applied FDR correction (*q* < 0.05) to control for multiple comparisons across the 400 cortical regions.

To probe other factors that might be associated with intra-individual variability in functional organization, we tested, for each participant, the Spearman-rank correlation between intra-individual daily variability in S-A axis loadings and 19 brain maps from the openly available Neuromaps database (https://github.com/netneurolab/neuromaps; [64]). We conducted a spin-permutation test with 1000 spherical rotations for each correlation analysis to control for spatial autocorrelation [104], and then applied FDR correction (*q* < 0.05) to control for multiple comparisons across the 19 tests conducted per subject. The following 19 brain maps were selected for their hypothesized relevance to intra-individual variability in S-A axis loadings: The first principal component of the 123 Neurosynth terms in the Cognitive Atlas, which represents meta-analytically-derived brain functions associated with cortical areas [105]; The first principal component computed for the top 1000 genes displaying the greatest variation in expression between cortical gyri of two brains recorded in the Allen Human Brain Atlas [65]; Metabolic measures such as glucose, oxygen, and cerebral blood flow [106]; Receptor densities of dopamine [107], acetylcholine [108], serotonin [109], norepinephrine [110], and glutamate [111]; Structural measures obtained from the Human Connectome Project S1200 release [112], including group average cortical myelin that was quantified using MRI T1-weighted/T2-weighted ratio [113] and cortical thickness; Electrophysiological MEG power distributions from 6 frequency bands, also obtained from the Human Connectome Project S1200 release [112], including alpha (8-12 Hz), beta (15-29 Hz), delta (2-4 Hz), low gamma (30-59 Hz), high gamma (60-90 Hz), and theta (5-7 Hz); And a representation of evolutionary expansion, based on the cortical surface area expansion from macaque to human [114].

We used linear regression to investigate local effects of hormone levels and perceived stress on the S-A axis loadings in two sets of analyses. In our first set of analyses, estradiol and progesterone levels were included as covariates in the linear model testing hormonal effects in the female participant, and testosterone and cortisol levels were included as covariates in the linear model testing effects in the male participant. Separate linear models were used to account for the local effects of perceived stress (PSS score). We used FDR correction to control for multiple comparisons of the tested effects across the 400 cortical regions (*q* < 0.05).

To investigate system-level effects of steroid hormone levels and perceived stress on the S-A axis loadings, we used measures of network topology, which describe the physical organization of nodes in networks and of networks along the S-A axis. For this, we computed measures of within-and between-network dispersion, as described in previous work [60, 66]. Within-network dispersion is defined as the sum of the Euclidean distances squared between network nodes (represented by the parcel S-A axis loadings) to the network centroid (quantified by the median of S-A axis loadings for parcels belonging to the same network), for which a higher value indicates a wider distribution of a given network’s nodes along the S-A axis, indicating greater segregation of the network. Between-network dispersion is defined as the Euclidean distance between network centroids, for which a higher value indicates that networks are more segregated from one another along the S-A axis. Within-network dispersion was computed for each of the seven Yeo-Krienen functional networks [67], and between-network dispersion was computed for each of the 21 possible network pairs. Then, to test for effects of hormone levels and perceived stress on measures of within-and between-network dispersion, we used the same linear models that we used to test for local effects. In order to assess statistical significance, we corrected for multiple comparisons, at Bonferroni-corrected thresholds of *p* < 0.004 (0.025/7) for the within-network effects and *p* < 0.001 (0.025/21) for the between network effects. For effects that survived Bonferroni correction, we further tested for their spatial specificity. Specifically, for each model, we generated a null distribution of *t*-values for the given effect using spin permutation testing (1000 spherical permutations) of the Schaefer 400 parcellation scheme, thus shuffling the network labels [104]. We thus controlled for spatial autocorrelation by assessing our empirical *t*-values against our generated null distributions, with a significance threshold of *p*_spin_ < 0.05.

Our second set of analyses aimed to test sex-specific effects of common steroid hormones (i.e., estradiol and testosterone) on functional organization in both participants. To do this, we replicated the analyses testing for local-and system-level effects described above, here including estradiol and testosterone as covariates in models tested in both participants.

## Author contributions

Conceptualization of original study design and data collection: LP, HG, and EGJ. Conceptualization of the current study and analyses: BS and SLV. Main analyses and visualizations: BS and DY. Writing—original draft: BS and DY. Writing— review and editing: BS, DY, LP, HG, EGJ, SBE, SLV. Supervision: SLV.

## Data availability

All data needed to evaluate the conclusions in the paper are present in the paper and the Supplementary Materials. We obtained data from the open-access 28&Me sample [46], available at https://openneuro.org/datasets/ds002674/versions/1.0.5, and the 28&He sample [47], soon to be available online.

## Code availability

The code used to conduct the analyses presented in this manuscript is available at https://github.com/biancaserio/MC_gradients. The code used to preprocess the data used in this manuscript is available at https://github.com/tsantander/PritschetSantander2020_NI_Hormones. The general code and tutorials for functional gradient decomposition can further be found at https://brainspace.readthedocs.io/en/latest/index.html.

## SUPPLEMENTARY INFORMATION

Supplementary results can be found in the Supplementary Materials.

