## Supplementary Material for "Exploring sex-specific neuroendocrine influences on the sensorimotor-association axis in single individuals"

† Shared first-author

* Correspondence to

Bianca Serio

Sofie L. Valk

**Supplementary Results**

**Figures**

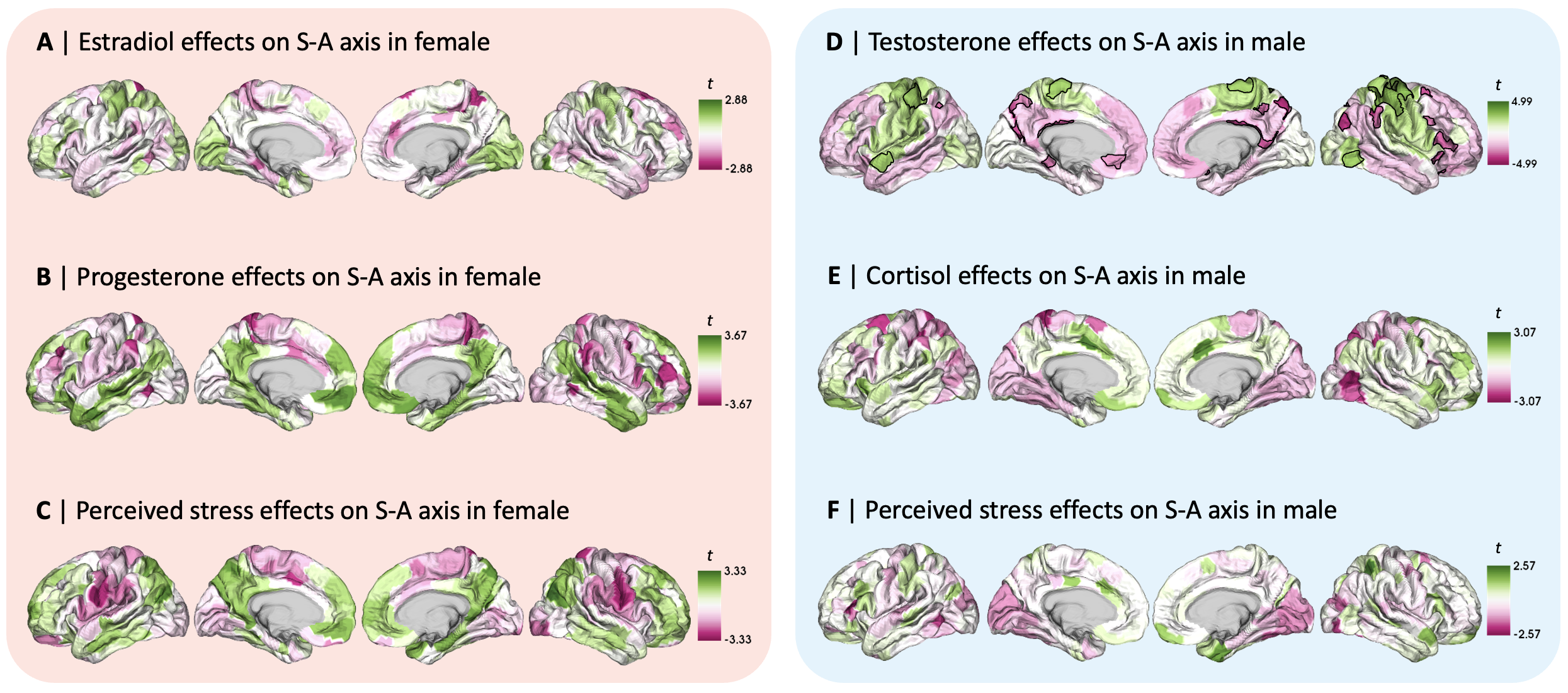

**Figure 1. Local-level effects on sensorimotor-association (S-A) axis loadings in the female and male participants. A** | Unthresholded *t*-maps of linear regression results showing patterns of local effects of **A** | estradiol, **B** | progesterone, and **C** | perceived stress on S-A axis loadings in the female participant; Unthresholded *t*-maps of linear regression results showing patterns of local effects of **D** | testosterone, **E** | cortisol, and **F** | perceived stress on S-A axis loadings in the male participant. Only the testosterone effects in the delineated cortical areas of panel D resulted as statistically significant following false discovery rate (FDR)-correction (*q* < 0.05).

**Tables**

|  | **Estradiol** | | | **Progesterone** | | |
| --- | --- | --- | --- | --- | --- | --- |
| **WN dispersion (network)** | ***t*** | ***p*** | ***p*_spin_** | ***t*** | ***p*** | ***p*_spin_** |
| Visual | 1.297 | 0.195 | - | -1.489 | 0.136 | - |
| Somatomotor | 1.296 | 0.195 | - | -0.577 | 0.564 | - |
| Dorsal attention | -0.085 | 0.932 | - | 0.439 | 0.660 | - |
| Ventral attention | -0.395 | 0.693 | - | -1.072 | 0.284 | - |
| Limbic | -0.196 | 0.844 | - | -0.620 | 0.535 | - |
| Fronto parietal | -0.621 | 0.535 | - | -2.625 | 0.009 | - |
| DMN | -0.696 | 0.486 | - | -1.299 | 0.194 | - |
| **BN dispersion  (pairwise networks)** | ***t*** | ***p*** | ***p*_spin_** | ***t*** | ***p*** | ***p*_spin_** |
| Visual - somatomotor | 0.299 | 0.765 | - | 0.430 | 0.667 | - |
| Visual - dorsal attention | 1.346 | 0.178 | - | -0.311 | 0.756 | - |
| Visual - ventral attention | 1.378 | 0.168 | - | 0.697 | 0.486 | - |
| Visual - limbic | 0.187 | 0.851 | - | -1.463 | 0.144 | - |
| Visual - fronto parietal | 0.382 | 0.702 | - | 0.258 | 0.796 | - |
| Visual - DMN | 0.866 | 0.387 | - | -1.656 | 0.098 | - |
| Somatomotor - dorsal attention | 0.539 | 0.590 | - | -0.644 | 0.520 | - |
| Somatomotor - ventral attention | 2.561 | 0.010 | - | 0.659 | 0.510 | - |
| Somatomotor - limbic | -0.132 | 0.895 | - | -1.424 | 0.155 | - |
| Somatomotor - fronto parietal | -0.007 | 0.994 | - | -0.194 | 0.846 | - |
| Somatomotor - DMN | 0.343 | 0.732 | - | -1.529 | 0.126 | - |
| Dorsal attention - ventral attention | 0.628 | 0.530 | - | 0.952 | 0.341 | - |
| Dorsal attention - limbic | -0.980 | 0.327 | - | -1.449 | 0.147 | - |
| Dorsal attention - fronto parietal | -0.822 | 0.411 | - | 0.604 | 0.546 | - |
| Dorsal attention - DMN | -0.187 | 0.852 | - | -1.720 | 0.085 | - |
| Ventral attention - limbic | -1.173 | 0.241 | - | -1.753 | 0.080 | - |
| Ventral attention - fronto parietal | -0.969 | 0.333 | - | -0.447 | 0.655 | - |
| Ventral attention - DMN | -0.650 | 0.515 | - | -1.838 | 0.066 | - |
| Limbic - fronto parietal | 0.321 | 0.748 | - | 3.106 | 0.002 | - |
| Limbic - DMN | 0.992 | 0.321 | - | -0.274 | 0.784 | - |
| Fronto parietal - DMN | 1.005 | 0.315 | - | -3.792 | 0.000* | 0.008* |

**Table 1. Steroid hormone effects on within-network (WN) and between-network (BN) dispersion in the female participant.** Results are yielded by a linear model including estradiol and progesterone as covariates. * indicates statistical significance after correcting for multiple comparisons, at Bonferroni-corrected thresholds of *p* < 0.004 (0.025/7) for the within-network effects and *p* < 0.001 (0.025/21) for the between network effects, followed by an additional correction for spatial autocorrelation via spin-permutation testing (1000 permutations), with *p*_spin_ < 0.05. DMN, default mode network.

|  | **Testosterone** | | | **Cortisol** | | |
| --- | --- | --- | --- | --- | --- | --- |
| **WN dispersion (network)** | ***t*** | ***p*** | ***p*_spin_** | ***t*** | ***p*** | ***p*_spin_** |
| Visual | 1.322 | 0.186 | - | 0.146 | 0.884 | - |
| Somatomotor | -0.738 | 0.460 | - | 0.943 | 0.346 | - |
| Dorsal attention | 2.943 | 0.003* | 0.005* | -0.627 | 0.530 | - |
| Ventral attention | 2.084 | 0.037 | - | 0.110 | 0.912 | - |
| Limbic | 1.078 | 0.281 | - | -0.340 | 0.734 | - |
| Fronto parietal | 1.488 | 0.137 | - | -0.732 | 0.464 | - |
| DMN | 1.184 | 0.236 | - | -0.572 | 0.567 | - |
| **BN dispersion  (pairwise networks)** | ***t*** | ***p*** | ***p*_spin_** | ***t*** | ***p*** | ***p*_spin_** |
| Visual - somatomotor | -2.861 | 0.004 | - | -1.446 | 0.148 | - |
| Visual - dorsal attention | -1.113 | 0.266 | - | 1.455 | 0.146 | - |
| Visual - ventral attention | -2.595 | 0.009 | - | -2.211 | 0.027 | - |
| Visual - limbic | 1.312 | 0.189 | - | -1.212 | 0.225 | - |
| Visual - fronto parietal | 1.903 | 0.057 | - | -0.777 | 0.437 | - |
| Visual - DMN | 1.446 | 0.148 | - | -0.815 | 0.415 | - |
| Somatomotor - dorsal attention | 0.923 | 0.356 | - | 2.030 | 0.042 | - |
| Somatomotor - ventral attention | -0.731 | 0.465 | - | -1.398 | 0.162 | - |
| Somatomotor - limbic | 2.107 | 0.035 | - | -0.580 | 0.562 | - |
| Somatomotor - fronto parietal | 2.880 | 0.004 | - | -0.024 | 0.980 | - |
| Somatomotor - DMN | 2.248 | 0.025 | - | -0.234 | 0.815 | - |
| Dorsal attention - ventral attention | -1.256 | 0.209 | - | -2.622 | 0.009 | - |
| Dorsal attention - limbic | 1.986 | 0.047 | - | -2.052 | 0.040 | - |
| Dorsal attention - fronto parietal | 3.084 | 0.002 | - | -2.016 | 0.044 | - |
| Dorsal attention - DMN | 2.052 | 0.040 | - | -1.564 | 0.118 | - |
| Ventral attention - limbic | 2.019 | 0.044 | - | 0.015 | 0.988 | - |
| Ventral attention - fronto parietal | 2.517 | 0.012 | - | 0.583 | 0.560 | - |
| Ventral attention - DMN | 2.085 | 0.037 | - | 0.300 | 0.764 | - |
| Limbic - fronto parietal | 0.210 | 0.833 | - | 1.497 | 0.134 | - |
| Limbic - DMN | 0.562 | 0.574 | - | 1.649 | 0.099 | - |
| Fronto parietal - DMN | 0.105 | 0.917 | - | -0.563 | 0.573 | - |

**Table 2. Steroid hormone effects on within-network (WN) and between-network (BN) dispersion in the male participant.** Results are yielded by a linear model including testosterone and cortisol as covariates. * indicates statistical significance after correcting for multiple comparisons, at Bonferroni-corrected thresholds of *p* < 0.004 (0.025/7) for the within-network effects and *p* < 0.001 (0.025/21) for the between network effects, followed by an additional correction for spatial autocorrelation via spin-permutation testing (1000 permutations), with *p*_spin_ < 0.05. DMN, default mode network.

|  | **PSS score (female)** | | | **PSS score (male)** | | |
| --- | --- | --- | --- | --- | --- | --- |
| **WN dispersion (network)** | ***t*** | ***p*** | ***p*_spin_** | ***t*** | ***p*** | ***p*_spin_** |
| Visual | -2.085 | 0.037 | - | -1.011 | 0.312 | - |
| Somatomotor | -0.143 | 0.886 | - | -1.081 | 0.280 | - |
| Dorsal attention | -1.627 | 0.104 | - | 0.346 | 0.730 | - |
| Ventral attention | -1.190 | 0.234 | - | -0.500 | 0.617 | - |
| Limbic | -0.592 | 0.554 | - | -0.157 | 0.875 | - |
| Fronto parietal | -0.969 | 0.332 | - | -0.121 | 0.904 | - |
| DMN | -1.945 | 0.052 | - | -0.103 | 0.918 | - |
| **BN dispersion  (pairwise networks)** | ***t*** | ***p*** | ***p*_spin_** | ***t*** | ***p*** | ***p*_spin_** |
| Visual - somatomotor | 0.713 | 0.476 | - | 0.043 | 0.966 | - |
| Visual - dorsal attention | -2.168 | 0.030 | - | -1.964 | 0.050 | - |
| Visual - ventral attention | 0.562 | 0.574 | - | -0.625 | 0.532 | - |
| Visual - limbic | -1.780 | 0.075 | - | -0.857 | 0.391 | - |
| Visual - fronto parietal | -1.696 | 0.090 | - | -0.541 | 0.588 | - |
| Visual - DMN | -1.622 | 0.105 | - | -0.456 | 0.648 | - |
| Somatomotor - dorsal attention | -2.193 | 0.028 | - | -1.445 | 0.148 | - |
| Somatomotor - ventral attention | -0.221 | 0.825 | - | -0.903 | 0.367 | - |
| Somatomotor - limbic | -1.922 | 0.055 | - | -0.737 | 0.461 | - |
| Somatomotor - fronto parietal | -1.698 | 0.090 | - | -0.442 | 0.659 | - |
| Somatomotor - DMN | -1.780 | 0.075 | - | -0.398 | 0.691 | - |
| Dorsal attention - ventral attention | 2.038 | 0.042 | - | 0.550 | 0.582 | - |
| Dorsal attention - limbic | -0.209 | 0.834 | - | 0.006 | 0.995 | - |
| Dorsal attention - fronto parietal | 0.037 | 0.970 | - | 0.575 | 0.565 | - |
| Dorsal attention - DMN | -0.080 | 0.936 | - | 0.383 | 0.702 | - |
| Ventral attention - limbic | -1.797 | 0.072 | - | -0.316 | 0.752 | - |
| Ventral attention - fronto parietal | -1.611 | 0.107 | - | -0.003 | 0.998 | - |
| Ventral attention - DMN | -1.676 | 0.094 | - | -0.033 | 0.974 | - |
| Limbic - fronto parietal | 0.331 | 0.741 | - | 1.078 | 0.281 | - |
| Limbic - DMN | 0.167 | 0.867 | - | 1.784 | 0.074 | - |
| Fronto parietal - DMN | -0.163 | 0.870 | - | -0.107 | 0.915 | - |

**Table 3. Perceived stress effects on within-network (WN) and between-network (BN) dispersion in the female and male participants.** Results are yielded by independent linear models for each participant, which only include perceived stress scale (PSS) score as a covariate. None of the tested effects were statistically significant after correction for multiple comparisons, i.e., at Bonferroni-corrected thresholds of *p* < 0.004 (0.025/7) for the within-network effects and *p* < 0.001 (0.025/21) for the between network effects. DMN, default mode network.

|  | **Estradiol (female)** | | | **Estradiol (male)** | | |
| --- | --- | --- | --- | --- | --- | --- |
| **WN dispersion (network)** | ***t*** | ***p*** | ***p*_spin_** | ***t*** | ***p*** | ***p*_spin_** |
| Visual | 1.137 | 0.255 | - | -1.154 | 0.248 | - |
| Somatomotor | 1.223 | 0.221 | - | -1.478 | 0.139 | - |
| Dorsal attention | -0.034 | 0.973 | - | 0.123 | 0.902 | - |
| Ventral attention | -0.547 | 0.585 | - | -0.317 | 0.751 | - |
| Limbic | -0.305 | 0.760 | - | 0.082 | 0.934 | - |
| Fronto parietal | -0.954 | 0.340 | - | 0.394 | 0.694 | - |
| DMN | -0.927 | 0.354 | - | 0.279 | 0.780 | - |
| **BN dispersion  (pairwise networks)** | ***t*** | ***p*** | ***p*_spin_** | ***t*** | ***p*** | ***p*_spin_** |
| Visual - somatomotor | 0.367 | 0.714 | - | 0.463 | 0.643 | - |
| Visual - dorsal attention | 1.464 | 0.143 | - | -2.259 | 0.024 | - |
| Visual - ventral attention | 1.478 | 0.139 | - | -0.108 | 0.914 | - |
| Visual - limbic | -0.060 | 0.952 | - | -0.157 | 0.875 | - |
| Visual - fronto parietal | 0.415 | 0.678 | - | -0.213 | 0.832 | - |
| Visual - DMN | 0.615 | 0.539 | - | -0.098 | 0.922 | - |
| Somatomotor - dorsal attention | 0.443 | 0.658 | - | -1.903 | 0.057 | - |
| Somatomotor - ventral attention | 2.707 | 0.007 | - | -0.607 | 0.544 | - |
| Somatomotor - limbic | -0.369 | 0.712 | - | -0.338 | 0.735 | - |
| Somatomotor - fronto parietal | -0.054 | 0.957 | - | -0.419 | 0.676 | - |
| Somatomotor - DMN | 0.093 | 0.926 | - | -0.268 | 0.789 | - |
| Dorsal attention - ventral attention | 0.811 | 0.418 | - | 0.983 | 0.326 | - |
| Dorsal attention - limbic | -1.156 | 0.248 | - | 0.773 | 0.440 | - |
| Dorsal attention - fronto parietal | -0.731 | 0.464 | - | 0.984 | 0.325 | - |
| Dorsal attention - DMN | -0.423 | 0.672 | - | 0.727 | 0.467 | - |
| Ventral attention - limbic | -1.410 | 0.159 | - | -0.046 | 0.963 | - |
| Ventral attention - fronto parietal | -1.060 | 0.289 | - | -0.056 | 0.955 | - |
| Ventral attention - DMN | -0.918 | 0.359 | - | -0.012 | 0.990 | - |
| Limbic - fronto parietal | 0.692 | 0.489 | - | -0.015 | 0.988 | - |
| Limbic - DMN | 0.957 | 0.339 | - | 0.168 | 0.867 | - |
| Fronto parietal - DMN | 0.357 | 0.721 | - | 0.114 | 0.909 | - |

**Table 4. Estradiol effects on within-network (WN) and between-network (BN) dispersion in the female and male participants.** Results are yielded by a linear model including estradiol and testosterone as covariates. None of the tested effects were statistically significant after correction for multiple comparisons, i.e., at Bonferroni-corrected thresholds of *p* < 0.004 (0.025/7) for the within-network effects and *p* < 0.001 (0.025/21) for the between network effects. DMN, default mode network.

|  | **Testosterone (female)** | | | **Testosterone (male)** | | |
| --- | --- | --- | --- | --- | --- | --- |
| **WN dispersion (network)** | ***t*** | ***p*** | ***p*_spin_** | ***t*** | ***p*** | ***p*_spin_** |
| Visual | -2.699 | 0.007 | - | 0.907 | 0.365 | - |
| Somatomotor | -1.004 | 0.315 | - | 0.661 | 0.509 | - |
| Dorsal attention | -0.709 | 0.478 | - | 0.122 | 0.903 | - |
| Ventral attention | -0.074 | 0.941 | - | 0.057 | 0.954 | - |
| Limbic | -0.788 | 0.431 | - | -0.227 | 0.820 | - |
| Fronto parietal | -1.307 | 0.191 | - | -0.657 | 0.511 | - |
| DMN | -1.440 | 0.150 | - | -0.603 | 0.547 | - |
| **BN dispersion  (pairwise networks)** | ***t*** | ***p*** | ***p*_spin_** | ***t*** | ***p*** | ***p*_spin_** |
| Visual - somatomotor | 0.156 | 0.876 | - | -1.121 | 0.262 | - |
| Visual - dorsal attention | -2.961 | 0.003 | - | 0.984 | 0.325 | - |
| Visual - ventral attention | -0.317 | 0.752 | - | -0.108 | 0.914 | - |
| Visual - limbic | -1.861 | 0.063 | - | -0.335 | 0.738 | - |
| Visual - fronto parietal | -1.421 | 0.155 | - | 0.014 | 0.989 | - |
| Visual - DMN | -2.200 | 0.028 | - | -0.212 | 0.832 | - |
| Somatomotor - dorsal attention | -1.893 | 0.058 | - | 1.578 | 0.115 | - |
| Somatomotor - ventral attention | -1.119 | 0.263 | - | 0.898 | 0.369 | - |
| Somatomotor - limbic | -1.439 | 0.150 | - | 0.198 | 0.843 | - |
| Somatomotor - fronto parietal | -1.019 | 0.308 | - | 0.606 | 0.544 | - |
| Somatomotor - DMN | -1.628 | 0.104 | - | 0.257 | 0.797 | - |
| Dorsal attention - ventral attention | 1.320 | 0.187 | - | -0.574 | 0.566 | - |
| Dorsal attention - limbic | 0.144 | 0.886 | - | -0.734 | 0.463 | - |
| Dorsal attention - fronto parietal | 0.818 | 0.414 | - | -0.506 | 0.613 | - |
| Dorsal attention - DMN | -0.199 | 0.843 | - | -0.564 | 0.573 | - |
| Ventral attention - limbic | -1.017 | 0.309 | - | -0.165 | 0.869 | - |
| Ventral attention - fronto parietal | -0.621 | 0.535 | - | 0.074 | 0.941 | - |
| Ventral attention - DMN | -1.198 | 0.231 | - | -0.089 | 0.929 | - |
| Limbic - fronto parietal | 0.826 | 0.409 | - | 0.818 | 0.414 | - |
| Limbic - DMN | -0.434 | 0.664 | - | 0.348 | 0.728 | - |
| Fronto parietal - DMN | -1.359 | 0.174 | - | -0.517 | 0.605 | - |

**Table 5. Testosterone effects on within-network (WN) and between-network (BN) dispersion in the female and male participants.** Results are yielded by a linear model including estradiol and testosterone as covariates. None of the tested effects were statistically significant after correction for multiple comparisons, i.e., at Bonferroni-corrected thresholds of *p* < 0.004 (0.025/7) for the within-network effects and *p* < 0.001 (0.025/21) for the between network effects. DMN, default mode network.
